# Functionalist Oncology to Model the Contextuality of Dynamics and Treatment in Acute Myeloid Leukemia

**DOI:** 10.64898/2025.12.22.695906

**Authors:** Alexander Ehmann, Rakan Naboulsi, Martin Jädersten, Sylvain Tollis, Nikolas Herold

## Abstract

Acute myeloid leukemia (AML) is a disease with a high degree of intra- and interpatient heterogeneity. Current treatments almost uniformly employ the standardized ‘7+3’ regimen, which is based on cytarabine combined with an anthracycline, with long-term survival rates below 30%. Our work specifically focuses on the cytarabine component of this regimen. Functionalist oncology utilizes edge-weighted digraphs as a representational tool to model the mathematical functional interdependencies of disease factors. This methodology enables a conceptual and mechanistic analysis of key factors influencing therapeutic success, providing a framework that can be parameterized to explore patient-specific treatment responses when suitable data are available. It incorporates the oligoclonal nature of AML to perform, in principle, comprehensive cost-benefit analyses regarding the addition of individual drugs to existing treatment regimens and the number and type of chemotherapy courses to be given. Extrapolating from experimental data, we investigate the therapeutic potential and risks of SAMHD1 inhibitors in AML treatment as a proof-of-concept. We also provide a web-based interactive application to visualize hypothetical AML treatment scenarios, which can be combined with *ex vivo* single-cell phenotypic (gene expression), genotypic (somatic mutations), and functional (drug responses) analyses. Functionalist oncology can thus be used to generate testable hypotheses that might contribute to improving oncological decision-making, e.g., by identifying the optimal number, nature and sequence of chemotherapy blocks, including both existing AML-directed blocks and additional drugs, such as SAMHD1 inhibitors.

## Introduction

Systemic treatment with cytarabine (ara-C) in combination with an anthracycline is the backbone for intensive chemotherapy with curative intent of acute myeloid leukemia (AML) (DiNardo and Wei 2020). However, the 5-year overall survival rate in adult patients remains below 30% (Shallis, Wang et al. 2019), underscoring the need for more effective and personalized treatment strategies. Here, we present a proof-of-concept framework that aims to provide mechanistic insights with a focus on the ara-C/SAMHD1 axis within the ‘7+3’ induction regimen, which is given to patients, typically young adults, with curative intent.

AML is a complex, multifactorial disease whose progression and treatment response depend on the interplay of multiple contextual factors. Sterile α motif and HD domain-containing protein 1 (SAMHD1) plays a dual role in the dynamics of cancer, particularly in AML (Herold, Rudd et al. 2017, Schaller and Herold 2020). As a tumor suppressor (De Silva, Hoy et al. 2013, De Silva, Wang et al. 2013, Clifford, Louis et al. 2014, Wang, Lu et al. 2014, Rentoft, Lindell et al. 2016, Herold, Rudd et al. 2017, Coquel, Silva et al. 2018, Johansson, Klein-Hitpass et al. 2018, Kodigepalli, Bonifati et al. 2018, Schaller and Herold 2020, Schott, Majer et al. 2022), it maintains dNTP pool homeostasis required for DNA replication, regulates replication stress (Franzolin, Pontarin et al. 2013), and activates apoptotic pathways following DNA double-strand breaks, thereby reducing the proliferation of AML cells (Clifford, Louis et al. 2014, Daddacha, Koyen et al. 2017, Coquel, Silva et al. 2018). However, SAMHD1 also acts as an ara-CTPase, hydrolyzing the active metabolite of ara-C (ara-CTP) and reducing its therapeutic efficacy (Herold, Rudd et al. 2017, Herold, Rudd et al. 2017, Herold, Rudd et al. 2017, Hollenbaugh, Shelton et al. 2017, Rudd, Schaller et al. 2017). Notably, low expression of SAMHD1 correlates with significantly longer event-free and overall survival of AML patients treated with ara-C (Herold, Rudd et al. 2017, Rassidakis, Herold et al. 2018). Targeting SAMHD1 thus creates a therapeutic dilemma: SAMHD1 acts as a suppressor of AML but becomes a resistance factor during ara-C treatment by reducing its efficacy. This context-dependence highlights the need for a systematic approach to evaluate how factors like SAMHD1 influence treatment outcomes. The purpose of this work is to provide a conceptual and mechanistic proof of concept for incorporating contextual factors into the design of AML treatment strategies, supported by theoretical models and experimental tools.

In our framework, factors are categorized based on their direct or indirect effect on AML. While the lowest level (primary) factors, such as ara-C, usually have a well-defined, direct effect on AML (e.g., cytotoxicity), secondary (non-primary) factors, such as SAMHD1, affect AML indirectly through interacting with primary factors. Ara-C can be considered a favorable primary contextual factor due to its direct cytotoxic effect on AML cells. Hence, SAMHD1 is an unfavorable secondary contextual factor for treating AML due to its inhibitory role on ara-C. Yet, as a tumor suppressor, SAMHD1 is also a favorable primary factor in AML. Therefore, the overall functional role of SAMHD1 is not trivial and is context-dependent. Consequently, the functional role of introducing an even higher-level tertiary contextual factor inhibiting SAMHD1 with the purpose of regaining ara-C’s cytotoxicity is not trivial either. To address the issue of context-dependent factors affecting AML, we employ “functionalist oncology”, where the chemical, biological, or pharmacological mechanisms are not described explicitly but rather at a phenomenological level. Factors are assembled into a graph that is analyzed and studied mathematically. This approach generates a hypothesis that requires clinical validation before it can be translated into practice. To mathematically predict the overall impact of the pharmacological inhibition of SAMHD1 on AML cells, we use edge-weighted digraphs, where factors are represented as vertices, and their interactions are represented as weighted edges (Methods). Digraphs enable the identification of important factors, simulate interventions, and predict outcomes. While mathematical modeling has previously been employed in oncology (Roeder and Glauche 2008, Stiehl, Ho et al. 2014, Fuentes-Garí, Misener et al. 2015, Bangsgaard, Andersen et al. 2020, Salehi, Kabeer et al. 2021), our functionalist approach is novel in its use of graphs to predict the benefits of adding new drugs (such as SAMHD1 inhibitors) to existing chemotherapy regimens. While SAMHD1 inhibitors are not part of standard-of-care as of today, a clinical trial evaluates this strategy in primary treatment (Jädersten, Lilienthal et al. 2022) and is in preparation as well for relapsed and refractory setting (Lilienthal, Tao et al. 2025). To make this framework accessible for pre-clinical studies, we developed a web-based interactive application that enables researchers and clinicians to model and visualize the predicted outcome of treatment scenarios, possibly providing guidance for personalized AML therapies.

## Methods

### Clinical context and modeling scope

The analyses presented in this manuscript reflect the intensive ‘7+3’ induction treatment in AML patients who can tolerate this regimen, with ara-C administered in combination with an anthracycline. In this model, the ara-C/SAMHD1/AML interactions are investigated in the context of combining SAMHD1 inhibitors or other drugs with ara-C. The term “ara-C” is therefore used as a shortcut for “ara-C-based treatment”. As a consequence, the model predictions (see below) are to be considered only in the context (drug combinations etc.) where model parametrization experiments are performed (mimicking the clinical context). Our model does not assume that *in vitro* viability assays accurately reflect leukemic stem cell eradication. Rather, these assays are used as proxies for relative clone-intrinsic properties, such as drug sensitivity or SAMHD1-mediated reduction of proliferation, which also influence stem-cell-containing subpopulations.

### Graphs

Graphs represent abstract structures consisting of factors (vertices) connected by edges (interactions). In this study, edge-weighted digraphs, i.e., directed graphs with weighted edges, representing the strength of interactions, are employed. An edge-weighted digraph *G* =( *V, E, f* ) consists of a set of vertices *V* ={*v_i_, v _j_*, …}, a set of edges *E* ={(*v_i_, v _j_* ) *,…*} (directed, from *v_i_* to *v _j_*), and a function *f*, assigning weights 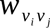 to each edge.

In this work, all edges represent inhibitory effects. Hence, weights range from 0 to 1 and represent probabilities or fractions of inhibition (e.g., the fraction of AML cells killed by a certain dose of ara-C). 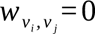 represents the absence of effect while 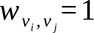 represents a total inhibitory effect.

In this framework, two inhibitory routes are modeled: Ara-C–mediated cytotoxicity and SAMHD1-mediated reduction of proliferation. Both routes were assumed statistically independent as a simplifying assumption in this proof-of-concept work. The statistical independence of the two routes of AML cell suppression must be understood as: 1) SAMHD1 does not reduce the activity of the ara-CTP metabolites that are not hydrolyzed, i.e., SAMHD1 neither interferes with ara-C cytotoxicity beyond ara-CTP hydrolysis, nor alters the downstream effect of ara-CTP such as DNA damage and repair. Reciprocally, the intrinsic anti-proliferative activity of SAMHD1 is not altered by the presence of ara-C; i.e., both in the presence and absence of ara-C, SAMHD1 expression reduces the proliferation of AML cells to the same extent. The model presented here applies as long as those assumptions are (at least approximately) valid.

### Analysis of Western blot and cell survival assays

To assess the impact of SAMHD1 on ara-C efficacy, we utilized data from Herold et al. (2017) (Herold, Rudd et al. 2017), which included Western blot analyses and cell survival assays. For the Western blot experiments, THP-1 cells were treated with varying amounts of virus-like particles (VLPs) containing Vpx, a protein known to induce SAMHD1 degradation via ubiquitination and proteasomal targeting (samples termed “X”), or empty VLPs (samples termed “dX”). SAMHD1 protein levels were quantified relative to the loading control GAPDH for each Vpx treatment, and band intensities were analyzed using Fiji software (Schindelin, Arganda-Carreras et al. 2012). For the cell survival assays, THP-1 cells were treated with increasing concentrations of ara-C in the absence or presence of varying amounts of Vpx-containing VLPs. Cell viability was measured after 72 hours. These *in vitro* measurements provide relative drug-sensitivity inputs and are not intended to represent patient-level dynamics. The cell viability data were analyzed across the different conditions to calculate model parameters as described in the Results. The uncertainty on model parameters was deduced from the uncertainty in viability measurements following rigorous error propagation procedures (Κυ 1966), under the assumption of statistically uncorrelated errors in distinct viability experiments. To quantify the experimental uncertainty in cell viability measurements, we calculated the coefficient of variation (CV) of viability for each ara-C concentration across samples with predicted wild-type SAMHD1 levels, including the dX and the no-VLP control samples (six samples in total). For each ara-C concentration, we calculated the mean and standard deviation of the viability values across those six samples, and the CV was computed as the ratio of the standard deviation to the mean (CV = SD/mean). This yielded 10 CV values (one per ara-C concentration), ranging from 0% to 23.8%, which we averaged to finally obtain an uncertainty of 9.4% in viability measurements (Supplementary Table 2).

### Software and data analysis

Graphs (Figs. 1, 2, 5) were created with Ipe 7.2.7, and the plot (Figure 7) was generated using Python 2.7.16 with Matplotlib 1.3.1 and NumPy 1.8.0. Phase diagrams (Figs. 3-4) were generated from the main text equations using custom scripts written in Matlab (Mathworks). Figure 6 was created using R (Team 2022) with the ggplot2 (Wickham, Chang et al. 2016) and Shiny (Chang, Cheng et al. 2022) packages.

**Figure 1.**
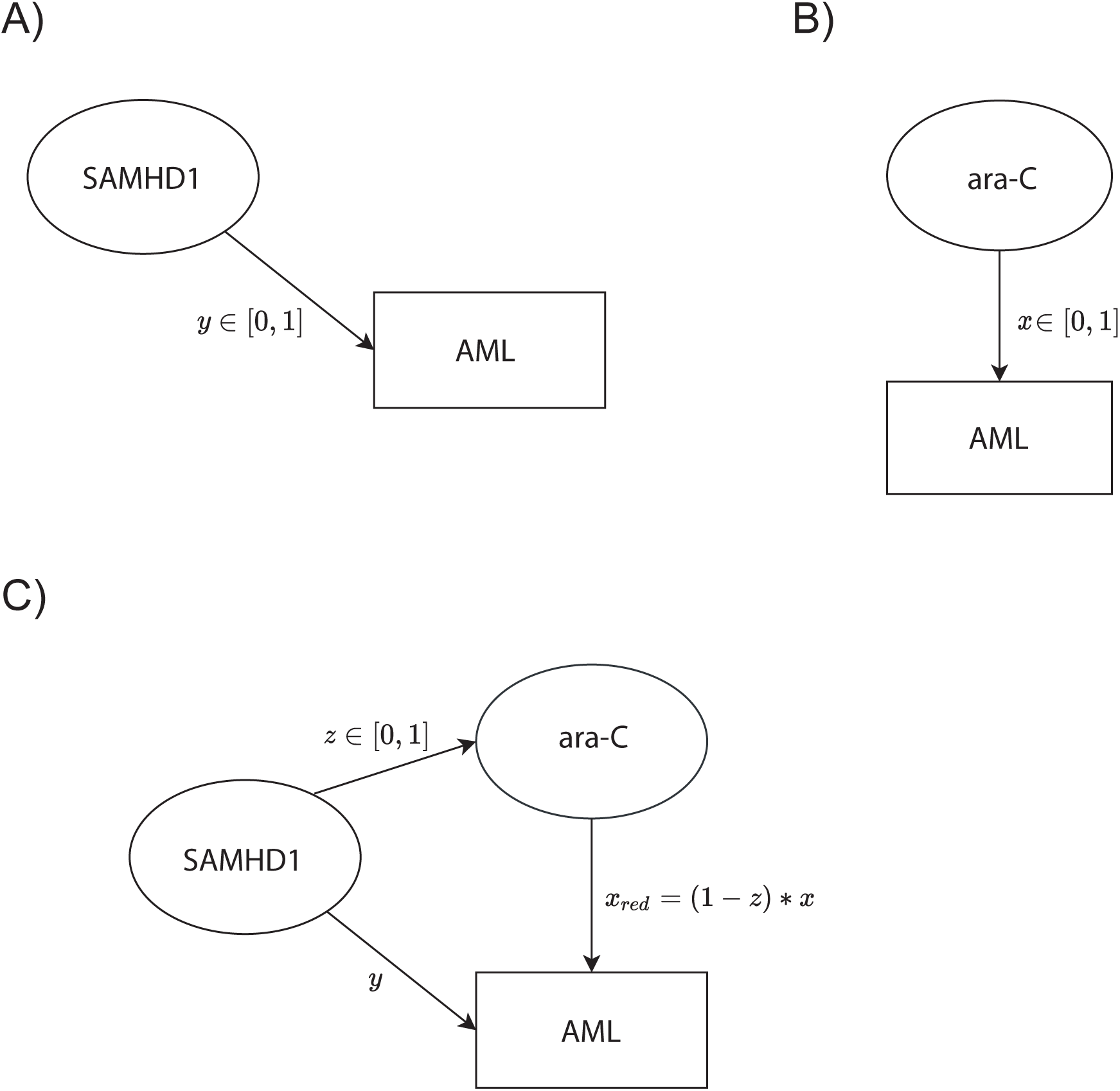
Digraph representation of monoclonal AML treatment with ara-C. (A) Digraph representing untreated monoclonal AML with the contextual factor SAMHD1 suppressing AML with efficacy y. (B) Digraph representing monoclonal AML treatment with a given dose of ara-C, which reduces the AML burden with efficacy x.(C) Digraph representing ara-C treated monoclonal AML with the contextual factors SAMHD1 also reducing the pool of active ara-C molecules, which, in turn, reduces ara-C efficacy towards AML by a factor z: *x_red_* =(1 - *z* ) * *x*.

**Figure 2.**
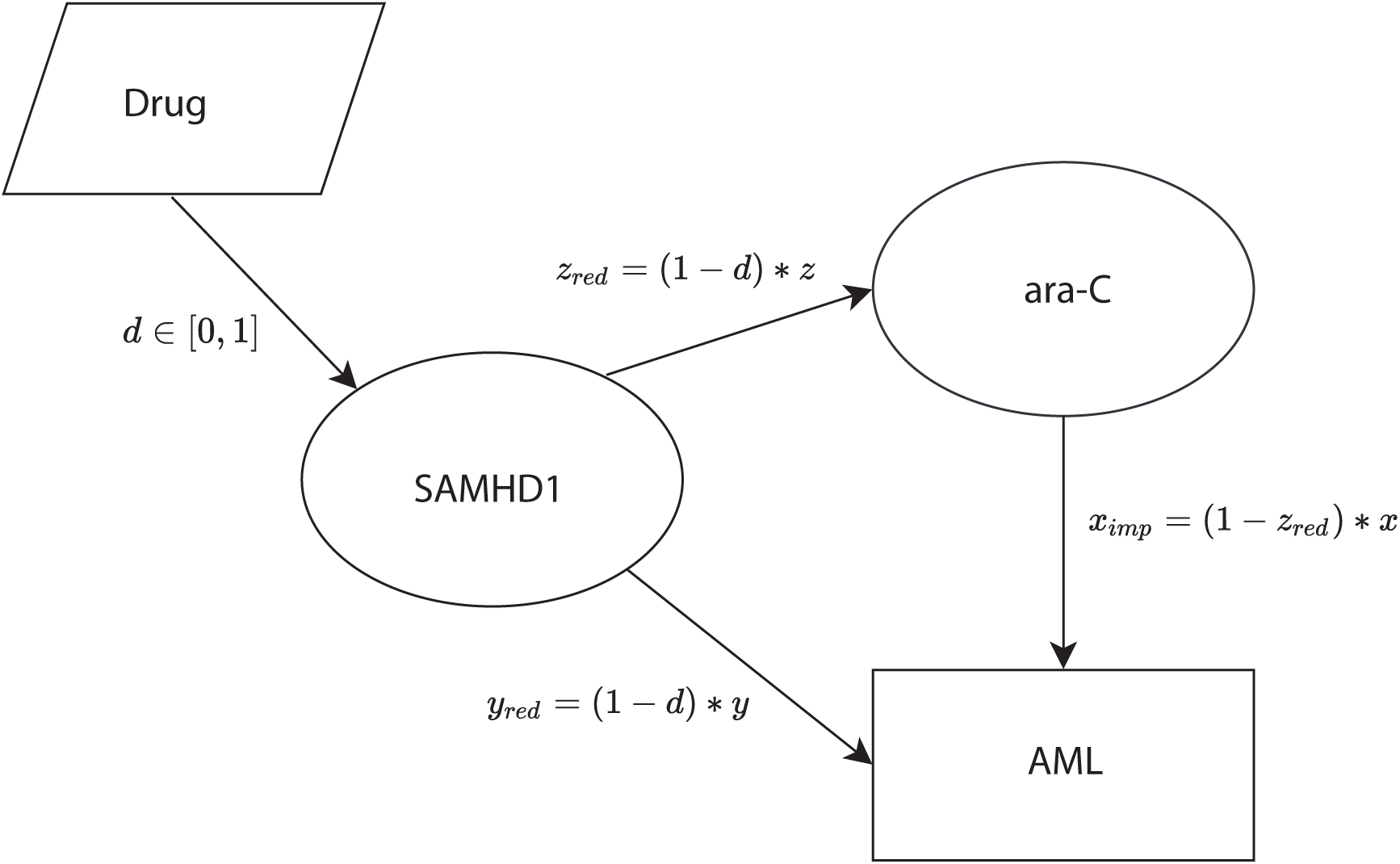
Digraph representation of monoclonal AML treatment with ara-C and anti-SAMHD1 drug. Efficacies of SAMHD1 towards AML and ara-C are reduced respectively to *y_red_* =(1 - *d* ) * *y*, *z_red_* =(1 - *d* ) * *z*, resulting in improved efficacy of ara-C towards AML *x_imp_* =(1 - *z_red_* )* *x*.

**Figure 3.**
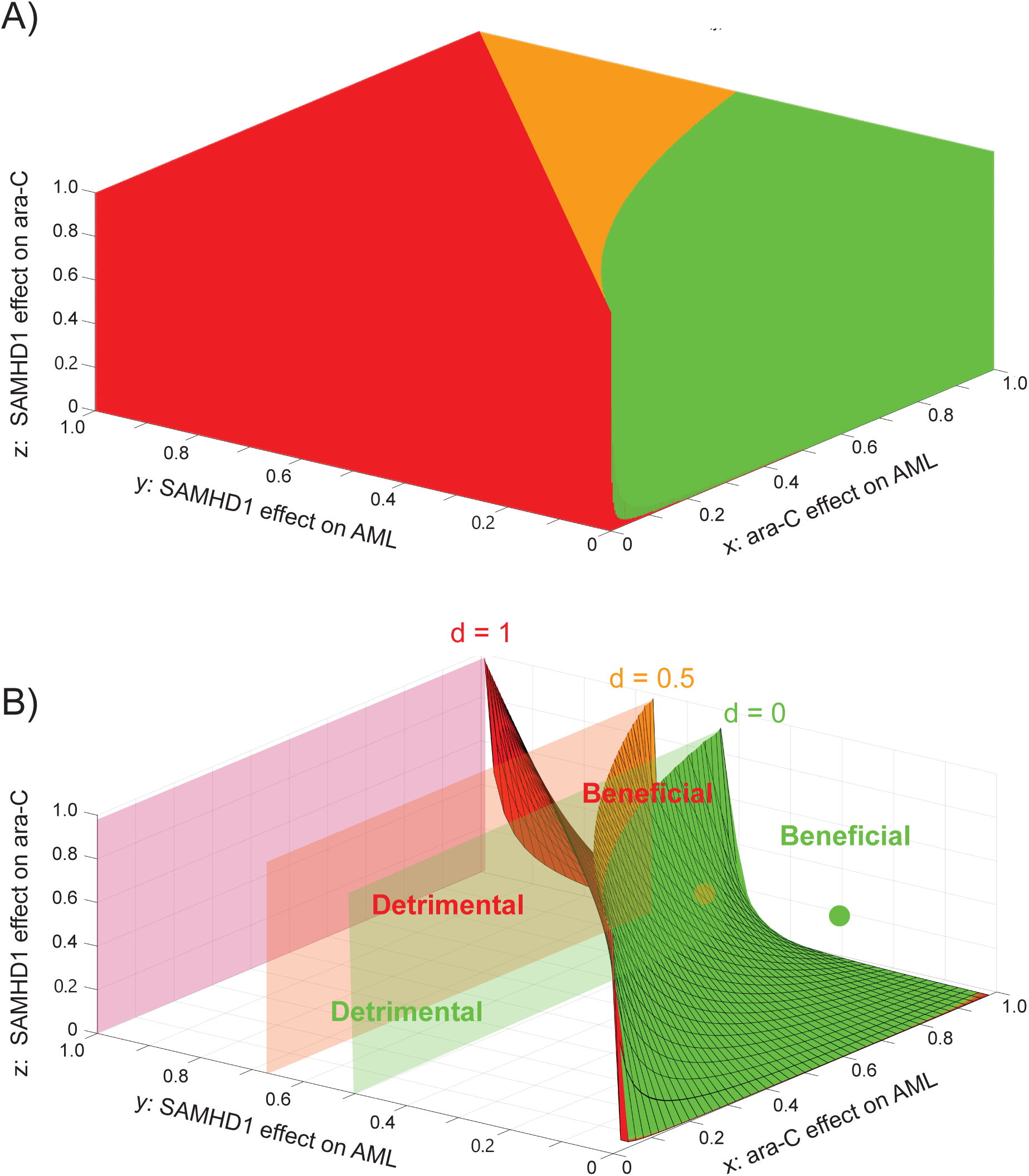
Mathematical analysis identifies quantitative conditions for a beneficial effect of targeting SAMHD1. A) State diagram of the AML/ara-C/SAMHD1/anti-SAMHD1 drug system, delineating regions of the (x,y,z) space where targeting SAMHD1 is always beneficial (green), always detrimental (red), or where benefits depend on the actual efficacy of the drug (orange). B) State diagram of the AML/ara-C/SAMHD1/anti-SAMHD1 drug system for specific values *d* = 0 (green surface), *d* = 0.5 (orange surface), and *d* = 1 (red surface). For each value of *d*, the (x,y,z) region below/above the corresponding limit surface corresponds to situations where targeting SAMHD1 is detrimental/beneficial, respectively. Specific examples of x, y and z discussed in the text are displayed as a green dot (situated above the green surface) and an orange dot (between the green and orange surfaces).

**Figure 4.**
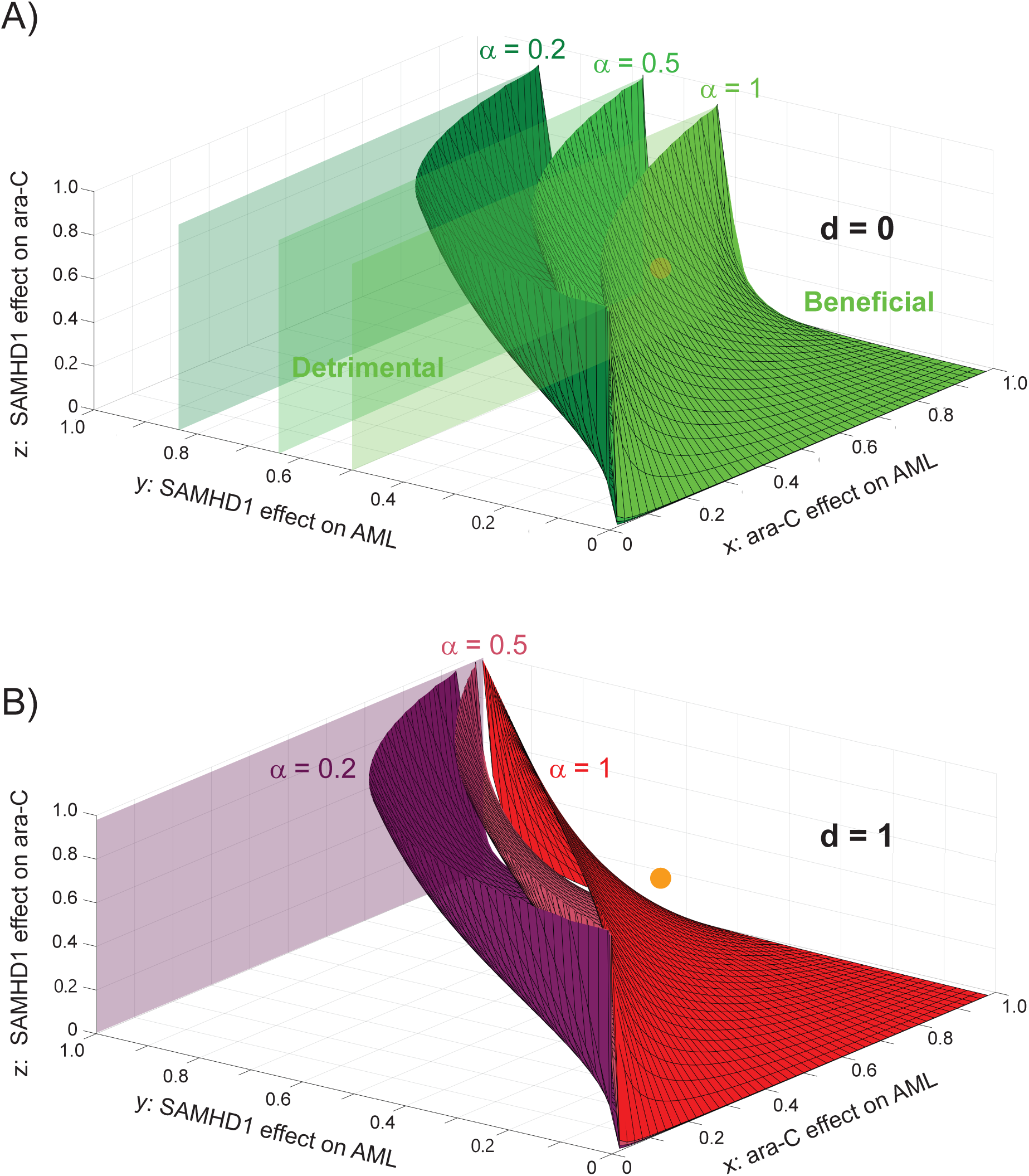
Mathematical analysis identifies quantitative conditions for a beneficial effect of specifically targeting SAMHD1’s ara-CTPase function. A) State diagram of the AML/ara-C/SAMHD1/anti-SAMHD1 drug system for d→0 (poorly effective drug), delineating regions of the (x,y,z) space where targeting SAMHD1 is beneficial for a poorly specific drug (α=1, light green), a moderately specific drug (α=0.5, medium green), and a highly specific drug (α=0,2, dark green). For each value of ɑ, the (x,y,z) region below/above the corresponding limit surface corresponds to situations where targeting SAMHD1 is detrimental/beneficial, respectively. B) State diagram of the AML/ara-C/SAMHD1/anti-SAMHD1 drug system for d=1 (very effective drug), delineating regions of the (x,y,z) space where targeting SAMHD1 is beneficial for a poorly specific drug (α=1, red), a moderately specific drug (α=0.5, dark pink), and a highly specific drug (α=0,2, violet). For each value of ɑ, the (x,y,z) region below/above the corresponding limit surface corresponds to situations where targeting SAMHD1 is detrimental/beneficial, respectively.

**Figure 5.**
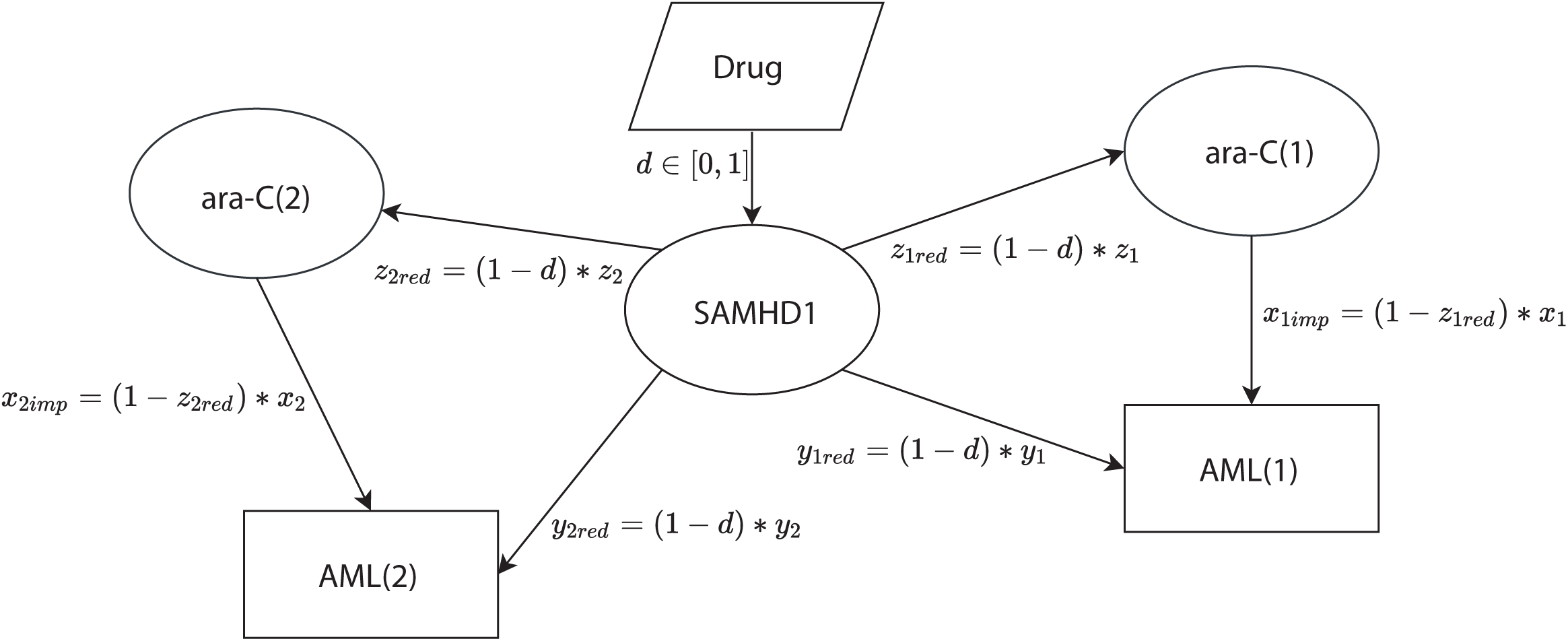
Digraph representation of oligoclonal AML treatment with ara-C and anti-SAMHD1 drug. The simple case of two clones is shown. Ara-C is represented as two nodes since the effects of SAMHD1 on ara-C and the effects of ara-C on the two clones AML (1) and AML (2) may be distinct. SAMHD1 is represented as only one node since it is assumed, on this digraph, that the effect of the drug on SAMHD1 is clone-independent.

**Figure 6.**
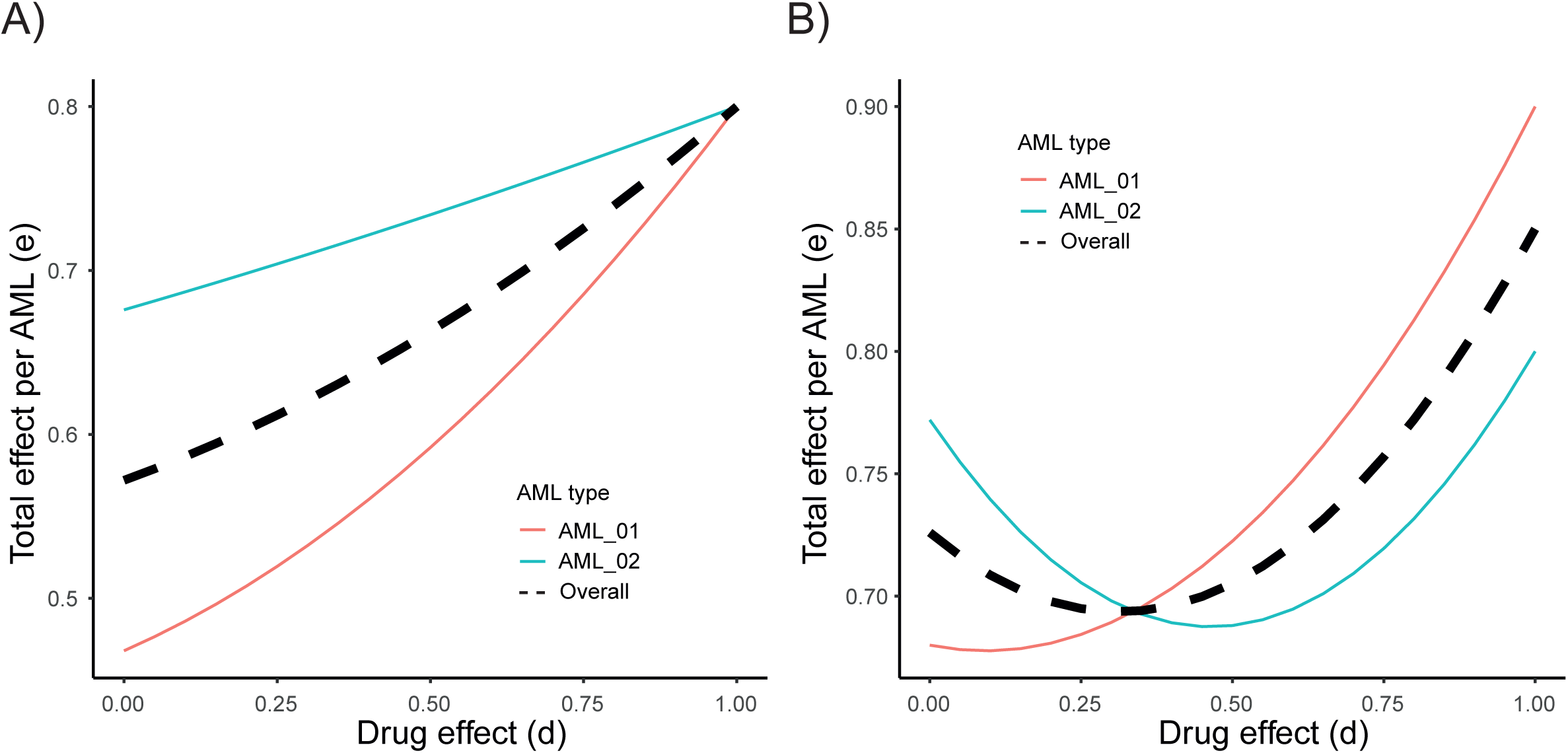
Oligoclonal AML treatment with ara-C and a SAMHD1 inhibitor. Total efficacy on the two AML clones AML(1) and AML(2), considered in equal fractions. Shown is the efficacy of the ara-C/SAMHD1 inhibitor combined treatment (y-axis) on AML(1) (red), AML(2) (blue) and overall (black dashed), as a function of the anti-SAMHD1 drug efficacy (x-axis) in the cases where SAMHD1 inhibition is beneficial for both clones irrespective of d (Left, x1=0.8, y1=0.3, z1=0.7, x2=0.8, y2=0.1, z2=0.2) and where SAMHD1 inhibition is beneficial only above a certain critical d value, different for both clones (Right, x1=0.9, y1=0.5, z1=0.6, x2=0.8, y2=0.7, z2=0.7).

**Figure 7.**
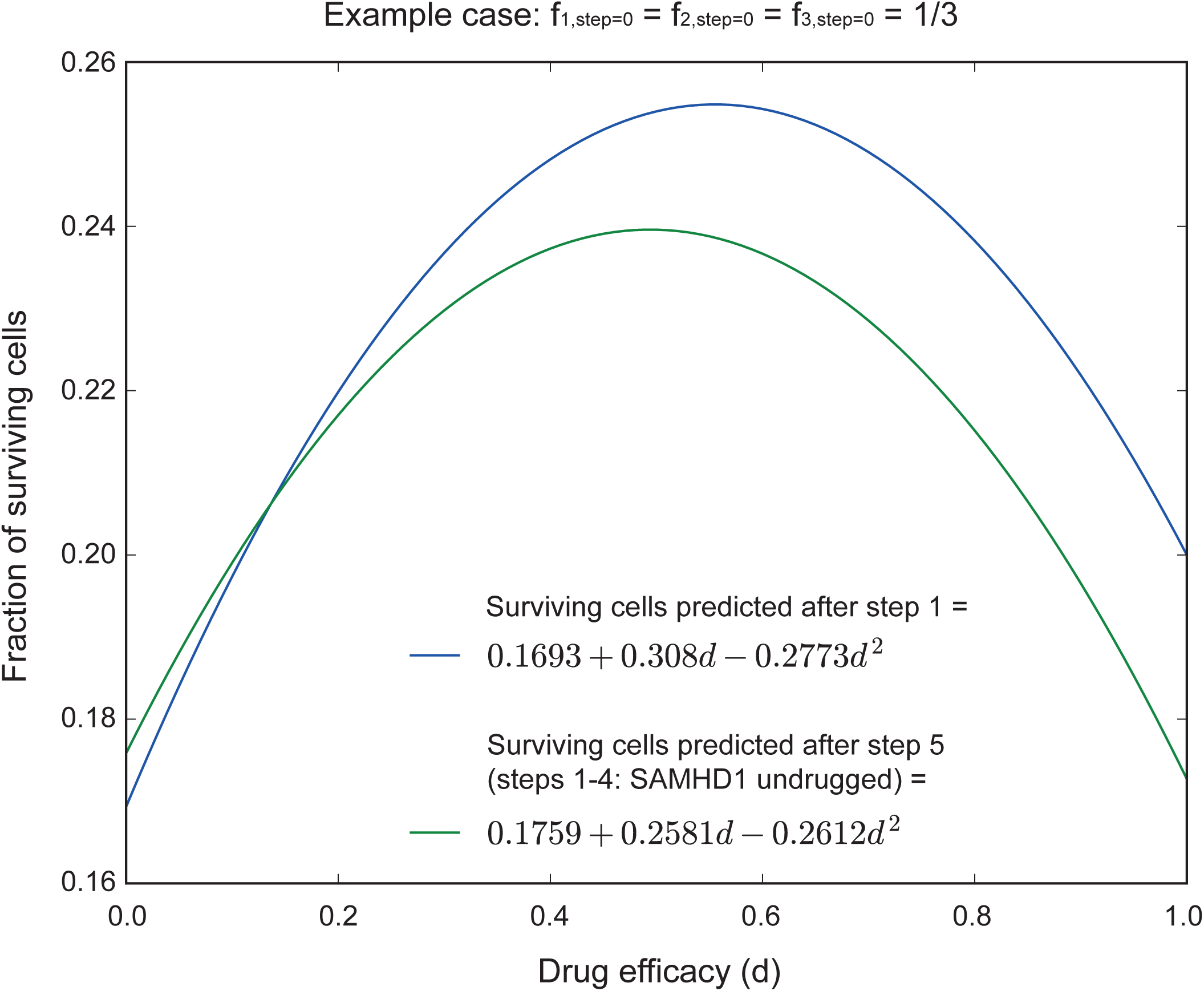
Sequential treatment of oligoclonal AML with ara-C and anti-SAMHD1 drug. Remaining AML burden (y-axis, AML cell population remaining after treatment in units of the expected population in the absence of treatment), predicted by the sequential model in the case of one cycle of treatment (blue) and five cycles of treatment (4 without drug and the fifth one with drug, green), as a function of the anti-SAMHD1 drug efficacy d.

## Results

### Modeling monoclonal AML treatment with ara-C

To address the issues related to the context-dependent effects of SAMHD1 and ara-C on AML, we developed a functionalist oncological mathematical model linking the factors ara-C, SAMHD1 and AML via a probabilistic description of their interactions.

SAMHD1 acts as a primary inhibitory factor on AML. We modeled this effect using a directed graph (digraph) consisting of a single direct edge from SAMHD1 to AML with weight *y ∈* [0,1] that represents the probability that an AML cell is not found in a population due to SAMHD1 (Fig. 1A). The magnitude of *y* depends on SAMHD1 expression levels (Herold, Rudd et al. 2017)^16,20^ . Similarly, we modeled the effect of ara-C on AML in the absence of SAMHD1 (Fig. 1B). Here, the edge weight *x ∈* [0,1] of the digraph represents the toxicity of ara-C on AML, interpreted as the fraction of AML cells killed by ara-C. The efficacy of ara-C towards AML cells is reduced by a fraction *z* by SAMHD1 (Forey, Cros-Perrial et al. 2021), which acts as a secondary disinhibitory factor (see Fig. 1C). The (reduced) efficacy of ara-C on AML is therefore *x_red_* =(1 - *z* ) * *x*. We stress that this equation defines z as one minus the ratio of ara-C cytotoxicity in the presence and absence of a given level/dose of SAMHD1. Hence, z depends on the expression levels of SAMHD1 and on ara-C doses, capturing the full enzymatic dynamics, and it is therefore not necessarily proportional to the fraction of ara-CTP molecules hydrolyzed by SAMHD1. In practice, probabilities may be identified with relative frequencies in large sample sizes. For instance, *y* is obtained by comparing the frequencies of surviving AML cells in the absence of ara-C and in the presence and absence of SAMHD1.

An AML cell survives the treatment if it is suppressed neither by the direct SAMHD1 route (Fig. 1A, with probability 1 - *y*) nor by the ara-C route (Fig.1B, with probability 1 - *x_red_*). In this work, both routes were assumed statistically independent. Under this assumption, AML cells survived with a probability (1 - *x_red_* )*(1 - *y* ) and thus the efficacy of the treatment could be expressed as one minus the fraction of surviving cells:

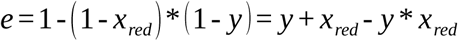

from which we obtained Equation 1:

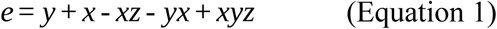

For example, if a certain dose of ara-C inhibits AML with efficacy *x* = 0.8 when unaffected by SAMHD1, and a certain amount of SAMHD1 inhibits ara-C with efficacy *z* = 0.7 and suppresses AML with efficacy *y* = 0.1, the resulting total treatment efficacy *e* = *y* + *x* - *xz* - *yx* + *xyz* = 0.316 was considerably lower than the inhibitory efficacy *x* = 0.8 of ara-C on AML without SAMHD1. This example demonstrated that SAMHD1 can be an unfavorable factor for treating AML with ara-C despite its functional role as a tumor suppressor. Therefore, SAMHD1 appears to be a crucial contextual factor that may need to be targeted, e.g., by introducing a tertiary contextual factor inhibiting SAMHD1 to regain the efficacy of ara-C towards AML.

### Modeling monoclonal AML treatment with ara-C and a SAMHD1 inhibitor

We first assumed the presence of a SAMHD1 inhibitor that affects equally the favorable (primary) tumor suppressor effect of SAMHD1 and the unfavorable (secondary) inhibitory effect on ara-C, both with the same efficacy *d ∈* [0,1] (Fig. 2). In this situation, *y* and *z* were reduced by a fraction *d* and became *y_red_* =(1 - *d* ) * *y* and *z_red_* =(1 - *d* ) * *z*, and as a consequence, the efficacy of ara-C towards AML *x_imp_* =(1 - *z_red_* )* *x* was improved compared to the case without a drug targeting SAMHD1. These calculations focus solely on the ara-C/SAMHD1 dynamics and do not account for other resistance pathways or co-administered drugs.

The efficacy of the combined treatment was directly derived from Equation 1 by substituting y and z with their reduced values (*e* = *y_red_* + *x* - *x z_red_* - *y_red_ x* + *x y_red_ z_red_*), leading to Equation 2:

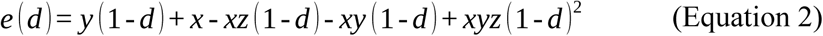

The introduced drug targeting SAMHD1 would be beneficial only if its efficacy in the presence of the drug is greater than in its absence. *e* ( *d* ) > *e* ( *d* = 0) is achieved if and only if *y* (1 - *d* ) + *x* - *xz* (1 - *d* ) - *xy* (1 - *d* ) + *xyz* (1 - *d* )^2^ > *y* + *x* - *xz* - *xy* + *xyz*, or equivalently if the condition below is satisfied:

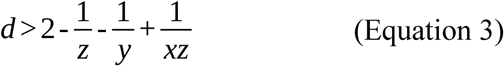

The drug would be detrimental otherwise. Two important conclusions emerged. First, irrespective of the drug efficacy, *d*, the inhibitory drug was always detrimental if *y* > *x*, i.e., if the direct effect of SAMHD1 on AML was stronger than that of fully active ara-C. Second, the inhibitor drug was always detrimental, regardless of *x* and *z*, if 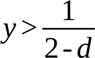. Hence, for any given efficacy of the SAMHD1-inhibitory drug, we identified an upper limit in *y* (the ara-C independent effect of SAMHD1 on AML) above which the drug was detrimental. Therefore, even though the balance between beneficial and detrimental roles of SAMHD1’s activity on AML involves, biologically, both ara-C dependent and independent pathways, there exists an ara-C independent condition (i.e., independent of *z* and *x*) on the net benefit of inhibiting SAMHD1.

Figure 3 illustrates the regions where the SAMHD1-inhibitory drug was beneficial (green), detrimental (red), or conditionally beneficial (orange). Increasing *x*, the efficacy of ara-C, or *z*, SAMHD1’s inhibitory effect on ara-C, generally increased the benefits of adding a SAMHD1 inhibitor, while increasing *y*, SAMHD1’s tumor-suppressive effect, produced the opposite outcome. For large enough *x* and *z* and small enough *y* (Fig. 3A, green region), the anti-SAMHD1 drug was overall beneficial.

Figure 3B shows the state space for *d=0* (green), *d=0.5* (orange), and *d=*1 (red). The drug was beneficial when *x, y,* and *z* defined a point situated above the surface corresponding to the value of *d*. Below these surfaces or when 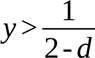, the drug was detrimental. Notably, the benefit *e* ( *d* ) - *e* ( *d* = 0), was proportional to *d*, therefore, small *d* yielded minimum gains or losses.

For example, in the case discussed above, where *x* = 0.8 *, y* = 0.1 and *z* = 0.7 (green dot in Fig. 3b), the drug was beneficial, irrespective of *d*. However, for other values like *x* = 0.7 *, y* = 0.4, *z* = 0.5, (orange dot), the inhibitory drug was only beneficial if *d* 0.3571.

Additional examples discussing the possible benefits of pharmacologically targeting SAMHD1 depending on the parameters *x*, *y* and *z*, which define distinct AML clones, are provided in Supplementary Figure 1a and b, where the total efficacy *e* ( *d* ) in the presence of a SAMHD1-inhibitory drug (Equation 2) was compared with the efficacy of the ara-C treatment alone (Equation 1) as a function of *d*.

### Modeling monoclonal AML treatment with ara-C and a SAMHD1 inhibitor with increased efficacy towards SAMHD1’s ara-CTPase function

SAMHD1’s dNTPase function, which requires a tetrameric configuration, may differ mechanistically from its tumor suppressor properties (Daddacha, Koyen et al. 2017, Herold, Rudd et al. 2017, Herold, Rudd et al. 2017, Coquel, Silva et al. 2018, Rudd, Tsesmetzis et al. 2020). This indicates that it is possible to design a SAMHD1 inhibitor that can preferentially target SAMHD1’s ara-CTPase function (*d_z_*) over its tumor suppressor function (*d _y_*). In this situation, the drug reduces *y* and *z* asymmetrically: *y_red_* =(1 - *d _y_* )* *y* and *z_red_* =(1 - *d_z_* )* *z*. Using Equation 1, we derived for the drug to be beneficial:

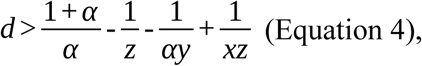

with *d* = *d_z_* and 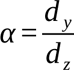

The case *α* = 1 describes the situation studied in the previous section, where the drug affects both functions equally. However, decreasing *α* (i.e., increasing specificity for the ara-CTPase function) moved the green and red surfaces in Fig. 3B leftwards (Fig. 4A, B), extending the space for which the drug is beneficial. Hence, our model predicted that it is always beneficial to specifically target the ara-CTPase function of SAMHD1 to improve treatment, compared to targeting both functions non-specifically. Increasing the drug specificity may be used as an alternative option to increasing drug efficacy in order to improve the overall treatment, for certain ranges of *x*, *y*, and *z* values predicted by our model (Fig. 4).

For example, in the case *x* = 0.7 *, y* = 0.4, *z* = 0.5(orange dot in Fig. 3b), when *d* is small *d*), a non-specific (*α* = 1) drug would be detrimental (Fig. 4A). However, increasing specificity (e.g., *α* = 0.5) shifted the orange dot into the beneficial region demonstrating that designing drugs that specifically target the ara-CTPase function of SAMHD1 (lower *α*), even at lower drug efficiencies (lower *d*), might improve treatment efficacy.

As in the case of Equation 3, we found an ara-C independent condition yielding detrimental effects: 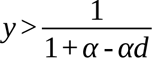. This condition is visualized in Fig. 4A (for *d* → 0) and Fig. 4B (for *d* = 1).

In summary, our model predicted that the treatment of AML with ara-C is not necessarily enhanced by supplementing with a SAMHD1-inhibitory drug. Such a drug should be administered only if it can significantly increase the total efficacy of the treatment as defined by the threshold in Equation 3. Otherwise, it might have a negative effect on the patient’s outcome. We identified two independent ways to improve efficacy: increasing the drug efficacy (Fig. 3B, green to red), a strategy constrained by the red curve ( *d* = 1), or increasing the drug specificity for SAMHD1’s ara-CTPase function (Fig. 4, light green/red to dark green/violet). The second approach was found to be more effective in the low *x* and *y* region (i.e., where both SAMHD1 and ara-C are poorly effective ). The best strategy to increase efficacy depends on the values of *x, y* and *z*, which we measured experimentally in the following section.

### Experimental determination of model parameters

To measure *x, y* and *z* parameters, we reanalyzed previously published data (Supp. Fig. 4A from (Herold, Rudd et al. 2017)). In these experiments, SAMHD1^+/+^ THP-1 AML cells were treated with ara-C and the simian immunodeficiency virus (SIV) protein Vpx, a protein that reduces SAMHD1’s protein levels by inducing its ubiquitination and proteasomal degradation (Schaller, Pollpeter et al. 2014) (Supplementary Table 1). The amounts of SAMHD1 in cells treated with increasing amounts of Vpx-containing Virion-Like Particles (VLPs, X) or empty VLPs (dX) relative to the no-VLP cells were estimated using GAPDH as a loading control (Methods). SAMHD1 levels decreased to ratios ranging from 0.82, with 40-fold diluted 2.5% VPX-containing VLPs, to as low as 0.006, with undiluted (100%) Vpx-containing VLPs (Supplementary Table 1). Hence, Vpx-containing VLPs constitute an experimental tool for controlling SAMHD1 expression.

Using Equation 1 to interpret the data, we analyzed cell viability under varying ara-C concentrations and SAMHD1 levels to estimate the parameters *x, y,* and *z*. Specifically, we used the following strategy.

First, we used the data with 100% Vpx-treated cells (where SAMHD1 levels are negligible) to estimate *x*, the dose-dependent effect of ara-C. In the absence of SAMHD1 ( *y* = *z* = 0), Equation 1 reduced to *e* = *x*, allowing *x* to be directly calculated as 1 - *viability*. These values are reported in Supplementary Table 1.

Then, we estimated *y*, the primary effect of SAMHD1 on AML, using data from cells in the absence of ara-C (*x=0*). Specifically, we compared the viability of cells treated with different amounts of Vpx with the average viability of the empty VLP (dX) and no-VLP controls. In this situation (*x=0*), *e* = *y* + *x* - *xz* - *yx* + *xyz* = *y*, so *y* was calculated as 1 - *viability*. Strikingly, despite the strong reduction in SAMHD1 levels when increasing the amount of Vpx (Supplementary Table 1), cell viability consistently remained at 100% compared to 90.18% in the six control samples treated with varying amounts of empty VLPs (dX) or untreated (no-VLP) (*P*=0.00172). Hence, reducing SAMHD1 beyond levels achieved with 2.5% Vpx (82% SAMHD1) does not further compromise cell viability. In short, we obtained *y=0.0982* for endogenous SAMHD1 levels, and *y=0* for all other (lower) SAMHD1 levels.

Finally, to measure *z*, the effect of SAMHD1 on ara-C, we used viability data of cells that have SAMHD1 at endogenous levels (where *y=0.0982*) treated with varying doses of ara-C (*x* obtained from Supplementary Table 1). Solving Equation 1 for *z*:

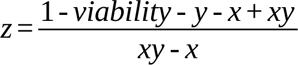

We found that SAMHD1 completely inhibits ara-C at concentrations up to 1.7 µM (z=1, up to experimental uncertainties, Supplementary Table 2). However, this inhibition decreases gradually with increasing ara-C concentration, dropping to *z* = 0.11 at 46 µM (clinically achievable concentrations).

In summary, this approach represents a functional assay to estimate *x*, *y*, and *z*, enabling us to use our model to practically estimate the benefits of targeting SAMHD1 in different cell samples.

### Modeling oligoclonal AML treatment with ara-C and a SAMHD1 inhibitor

AML is inherently heterogeneous, consisting of multiple clones and sub-clones that differ in their genotype and phenotype (Bochtler, Stölzel et al. 2013, Zachariadis, Cheng et al. 2020). The different clones of AML respond differently to the same therapy (Herold, Rudd et al. 2017, Morita, Wang et al. 2020), indicating that *x*, *y* and/or *z* depend on the AML clone. Although theoretically each AML cell may be considered as a single clone, in practice, a limited number of genetically distinct clones, or phenotypically distinct cell states (e.g., leukemic stem cells vs differentiated cells (Potter, Miraki-Moud et al. 2019, Velten, Story et al. 2021)) usually dominate an AML cell population (Ding, Ley et al. 2012, Walter, Shen et al. 2012), prompting us to adjust our framework to the case of oligoclonal AML with a discrete number of clones.

We modeled oligoclonal AML as a population of AML comprising *N* clones *C_i_* (where *i* = 1. . *N* ), each representing a fraction *f _i_* of the total cell population (_∑_ *f _i_* = 1). Clonal fractions *f _i_* can be determined by single-cell phenotypic and genotypic analyses (see “Limitations of Study and Outlook”) with three parameters characterizing each clone: *x_i_*, the effect of ara-C, *y_i_*, the tumor suppressive effect of SAMHD1, and *z_i_*, the inhibitory effect of SAMHD1 on ara-C. Moreover, *d_i_* represents the efficacy of the SAMHD1-inhibitory drug, which may vary across clones. To simplify, we only considered the symmetric case where the drug equally affects SAMHD1’s functions as an ara-CTPase and a tumor suppressor. However, the framework may be extended to asymmetric functions (*α_i_* < 1). The graph describing this situation is represented in Fig. 5 for two clones, with the drug affecting SAMHD1 equivalently across clones.

To compute the efficacy of treatment of oligoclonal AML, we defined *e* = 1 - *P* ( *survival* ), where *P* ( *survival* ) is the probability that a cell survives treatment. Using the law of total probability:

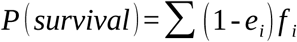

where *e_i_* is the efficacy towards clone *C_i_* (calculated using Equation 3), and *f _i_* is the fraction of cells in clone *C_i_*. From this, we get the total efficacy:

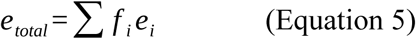

This assumes all clones are targeted equally. However, clinicians may prioritize certain clones by assigning weights *w_i_* (where _∑_ *w_i_* = 1) to reflect therapeutic goals. Although our model uses fixed clone-specific parameters, the framework allows these parameters to be updated between treatment cycles if biological changes (e.g., plasticity or microenvironmental adaptation) are suspected.

To illustrate Equation 5, we modeled a patient with two AML clones in equal proportions (*f* _1_ = *f* _2_ = 0.5). Clone AML(1) expresses SAMHD1 at a high level ( *y*_1_ = 0.3 and *z*_1_ = 0.7), making it less responsive to ara-C, while clone AML(2) expresses SAMHD1 at a low level ( *y*_2_ = 0.1 and *z*_2_ = 0.2). Both clones have similar intrinsic responses to ara-C (*x*_1_ = *x*_2_ = 0.8), and in both clones, SAMHD1 responds equally to the drug (*d*_1_ = *d*_2_ = *d*). In this case, the drug is beneficial to both clones regardless of *d*. Therefore, the drug is beneficial to oligoclonal AML, with a total efficacy that falls between the efficacy on both clones (Fig. 6, left).

We next consider the situation where each clone benefits from the anti-SAMHD1 drug only beyond a certain threshold of drug efficacy (orange region in Fig. 3A), but where these thresholds are different for the two clones. As the situation where a threshold drug efficacy exists to elicit benefits of SAMHD1 inhibition occupies a substantial region of the diagram in Fig. 3A (orange region), the likelihood that an AML patient would host two clones within this region is not negligible. For example, *x*_1_ = 0.9, *x*_2_ = 0.8, *y*_1_ = 0.5, *y*_2_ = 0.7, *z*_1_ = 0.6, *z*_2_ = 0.7, the threshold efficacies, above which the anti-SAMHD1 drug is beneficial, are *d*_1_ = 0.93 and *d*_2_ = 0.19. For *d* between 0.19 and 0.93, inhibiting SAMHD1 is detrimental to the treatment of AML(1) and favorable to that of AML(2). The total efficacy as a function of *d* (as defined by Equation 5 with *x_i_, y_i_* and *z_i_* defined above) is represented in Fig. 6, on the right side, and reveals the existence of a threshold of drug efficacy above which SAMHD1 inhibition is beneficial to the overall treatment of the oligoclonal AML, although for this value of the efficacy, the drug is detrimental to AML(2).

### Modeling optimal treatment sequences

In the therapeutic strategy defined above, the choice of a chemotherapy regimen (ara-C +/- a SAMHD1 inhibitory drug) will asymmetrically affect different clones, causing fractions to evolve over time, making it necessary to re-evaluate the efficacy as clonal fractions change.

If *K* cells are present before treatment, the population of clone *C_i_* becomes *K f _i_* (1 - *e_i_* ) after treatment. As a result, the updated fraction of the clone *C_i_* becomes

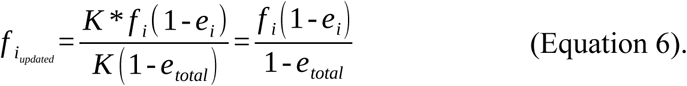

As a result, clones for which the efficacy *e_i_* > *e_total_* decrease in fraction, while those for which the efficacy *e_i_* < *e_total_* will increase. The updated total efficacy

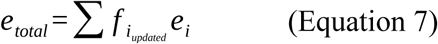

can then be analyzed to optimize the sequence of treatment strategies for individual patients.

For example, we assume a patient suffers from three AML clones in equal proportions 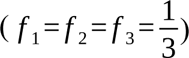 and parameters *y*_1_ = 0.6, = 0.6, *y*_2_, = 0.7, *y*_3_= 0.8, *z*_1_= 0.4, *z*_2_= 0.5, *z*_3_= 0.6, *x*_1_= 0.9, *x*_2_ = 0.8 and *x*_3_ = 0.7. Parameters were chosen to simulate a situation where, even after four cycles of ara-C treatment, AML is still not eradicated. AML treatment usually consists of two ara-C-containing induction courses followed by one up to three ara-C-containing consolidation courses and/or allogeneic haematopoietic stem cell transplantation (Jädersten, Lilienthal et al.2025). From Equation 5 we get 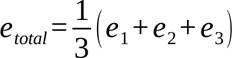, where *e_i_* = *x_i_* +(*y_i_* - *x_i_ z_i_* - *x_i_ y_i_* )(1 - *d* ) + *x_i_ y_i_ z_i_* (1 - *d* )^2^ (from Equation 3). With those parameters, SAMHD1 inhibition is beneficial to clone 1 but detrimental to clones 2 and 3, and also detrimental to the total population. Hence, in this situation, a treatment without SAMHD1 inhibition would be initiated. Using Equations 6 and 7 alternatively (with *d=0*), our model predicts that over a sequence of such ara-C only treatments, the fraction of clone 1 would increase in the overall population to the extent of the fractions of clones 2 and 3. The addition of a SAMHD1 inhibitor becomes favorable only after four cycles of ara-C treatment alone, provided the SAMHD1 inhibition is very strong (*d*→1, Fig. 7).

### What about in practice?

In practice, we suggest the following steps to evaluate our model predictions in pre-clinical test setups, with the long-term goal of bringing predictions into the clinic. Inspired by the analysis above, and adding an extra step to evaluate the efficacy of SAMHD1 functional inhibition from experimental data, the determination of a personalized treatment strategy would be:

- Run single-cell characterization and in vitro assays similar to Supplementary Tables 1-2, to identify the number of clones present in the patient, their fractions *f _i_* and the intrinsic response parameters *x_i_, y_i_* and *z_i_* of each clone.
- Maximize *e_total_* =∑ *f _i_ e_i_* ( *x_i_, y_i_, z_i_, d* ) with respect to the ara-C dose (which modulates *^x^_i_* but also affects *z_i_*), based on data similar to Supplementary Tables 1-2, and with respect to adding/not adding SAMHD1 inhibitory drug (of so far unknown efficacy *d*);
- Treat with ara-C +/- SAMHD1 inhibitory drug, re-measure the clonal fractions 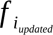 and fit the 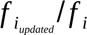 ratios for each clone to extract the drug efficacy *d* using as fitting formula

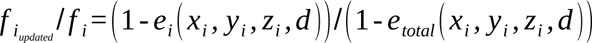
- Maximize 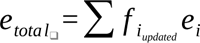 with respect to ara-C dose (which modulates *^x^_i_* but also affects *z_i_*), using tabulated values for those parameters, and for d.
- Treat with ara-C +/- SAMHD1 inhibitory drug
- Repeat the last two steps, maximizing 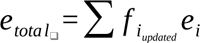 based on the updated fractions at each treatment phase.

### AsymSAM: a web-based interactive App for modeling oligoclonal AML

To facilitate the evaluation of various AML treatment scenarios involving multiple AML clones and a range of efficacy values, we have developed a web-based interactive application. This tool is designed to model and visualize the concurrent treatment efficacies of up to ten distinct AML clones. It is based on Equation 3 (to calculate the efficacy of a treatment towards each clone independently), Equation 5 (to calculate the overall efficacy of a treatment towards the whole AML cell population), and Equation 6 (to update the clonal fractions at each step of the treatment).

The application is designed to address the complexity of oligoclonal AML, where patients often have several different AML clones. It allows users to input and adjust efficacy values for three key parameters: the direct effect of ara-C on AML ( *x*), the direct effect of SAMHD1 on AML ( *y*), and the direct effect of SAMHD1 on ara-C ( *z*). These efficacy values can be modified incrementally, enabling precise sensitivity analyses to determine their impact on overall treatment efficacy.

Each parameter can be independently adjusted for each AML clone. This flexibility enables the generation of detailed plots that visualize both the total effect without SAMHD1 inhibition (*e_total_* (*d* = 0)) and the total effect with the drug (*e_total_* (*d* )) across all AML clones, either individually or overall, taking all clones into account.

## Discussion

The treatment of AML and other malignancies has evolved empirically through clinical trials. As of today, the single most effective drug combination to treat AML is ara-C with anthracyclines. Because anthracyclines, or any other co-administered agents, can alter ara-C exposure, leukemic cell states, and microenvironmental responses, clone eradication under combination therapy may exhibit non-independence. We and others have previously demonstrated that resistance to ara-C, mediated by resistance factors like SAMHD1, is a major reason for treatment failure. Accordingly, we have identified drugs with the potential to inhibit SAMHD1, which we are currently evaluating in a clinical trial (EUDRACT 2018-004050-16) (Rudd, Tsesmetzis et al. 2020, Herold 2021). However, AML is a complex disease with a heterogeneous population of leukemic cells. To cure AML, all malignant cells must be eradicated or transformed into a non-malignant state.

More generally, the applicability of the framework depends on the clinical context in which it is applied. The assumptions and parameters used here pertain only to the modeled ara-C/SAMHD1 axis under a specific therapeutic setting. In clinical combination regimens such as “7+3,” the probability of leukemic clone eradication may not be independent across agents, as anthracyclines or any other co-administered drug can influence ara-C pharmacokinetics, DNA damage response, or cell cycle distribution. Consequently, when the clinical context changes, all relevant model parameters must be re-estimated from experiments that explicitly capture the context. Future model extensions can incorporate such context dependence by adding interaction terms or conditional probabilities when combination-specific data become available. In the present work, we developed a functionalist approach to model treatment responses to ara-C in AML, consisting of different sub-entities (AML “clones”). We also modeled the role of SAMHD1 as a tumor suppressor and resistance factor for ara-C. This allowed a theoretical assessment of whether adding a SAMHD1 inhibitor would result in net beneficial effects, showing that the exact composition of a given AML largely determines the efficacy of a SAMHD1 inhibitor. We have further shown that a SAMHD1 inhibitor can have a negative effect on treatment outcomes. Hence, this theoretical framework allows for individualized treatment decisions. We have developed a web-based interactive application that enables modeling of scenarios with up to ten AML clones, supporting sample-based sensitivity analyses.

Evidently, the entire context is not always known. There might be relevant but unknown factors whose influence has not been defined. Other factors might be known to play a role but have not been identified. Yet other factors might be identified, but it is unclear which other factors they influence and how.

## Limitations of the Study and Outlook

What is currently lacking is a readily available empirical way to determine the AML architecture, expression levels of SAMHD1 in individual AML “clones,” relative efficacies of ara-C, and the inhibitory effects of a SAMHD1 inhibitor on individual AML “clones.” To this end, we are developing an experimental setup that compares the phenotypic and genotypic composition of AML patient blasts prior to and following exposure to single drugs and drug combinations *ex vivo*. We combine flow cytometry using clinically validated antibody panels (allowing for an immunophenotypic comparison of clonal composition) with, following single-cell sorting, DNTR-Seq (Zachariadis, Cheng et al. 2020). The latter combines scDNA-seq and scRNA-Seq simultaneously, allowing for an appreciation of genotypic and phenotypic properties. Hence, this setup allows an experimental determination of the number of AML “clones” and their proportions, expression levels of SAMHD1 in individual AML “clones,” empirical determination of the “weights” of ara-C inhibitory potential on different AML “clones”, as well as the change in number and proportion of AML “clones” following ara-C treatment with or without a SAMHD1 inhibitor. This procedure can be re-iterated, and even different treatment modalities can be used in sequence.

We also note that the efficacy equations (Equations 1-3) were calculated under the assumption of statistical independence between SAMHD1-mediated and ara-C-mediated suppression of AML cells. Strictly speaking, this assumption holds when a population of cells surviving ara-C is as likely as a population of cells sensitive to ara-C treatment to be affected by a given expression level of SAMHD1; and when, conversely, a population of cells surviving SAMHD1 expression is as likely as a population of cells sensitive to the same level of SAMHD1 expression to respond to ara-C treatment. Hence, genetic/epigenetic backgrounds that would influence both resistance/sensitivity to ara-C and resistance/sensitivity to SAMHD1 may require adjustments to the modeling framework to account for non-independent routes of ara-C and SAMHD1, which can be achieved mathematically through the use of conditional probabilities.

Our analysis of the viability data published by Herold and co-workers (Herold, Rudd et al. 2017) showed that, for THP-1 cells, reducing SAMHD1 levels by 18% was sufficient to increase viability to its maximum level. This result has to be compared with that of Kodigepalli et al. (Kodigepalli, Bonifati et al. 2018), where AML blasts were counted from patients’ blood samples and analyzed in regard to their SAMHD1 content, relative to THP-1 cells. Patients where SAMHD1 levels in blasts were very low showed 80% blast counts on average (normalized to the maximal blast count across all donors). In comparison, patients with SAMHD1 levels equal to 1-2.5 times that of THP-1 cells showed a median count of about 65%, with a high inter-donor variability. This corresponds to a SAMHD1-dependent viability reduction of (80-65)/65=18%, which is twice what we observed in our study. This difference might be explained by two non-exclusive factors: 1) in Kodigepalli et al. (Kodigepalli, Bonifati et al. 2018), data is missing between 50% and 100% SAMHD1 levels, i.e. the range in which most of the SAMHD1-related decrease in viability is expected, based on our results; hence, threshold effects may have been missed; and 2) Kodigepalli et al. (Kodigepalli, Bonifati et al. 2018) assessed SAMHD1 levels and viability levels in primary AML blasts, not in THP-1; primary AML and THP-1 cells may display different viability responses to SAMHD1 levels; for instance, AML cells from different donors in (Kodigepalli, Bonifati et al. 2018), show a broad range of viabilities although they have similar SAMHD1 levels, buttressing the fact that the parameters x, y, and z must be estimated for each particular cell population.

Our analysis of the case of oligoclonal AML is restricted to the case where the nature and the number of clones do not change throughout each step of the treatment and observation (only clonal fractions change). In the situation where new clones are detected in the course of the treatment, either due to de novo mutations or due to the positive selection pressure of the treatment on very rare, undetected clones in the initial state, those clones need to be experimentally assessed (to obtain the corresponding x, y and z values) before they can be included in the framework and a new treatment can be optimized. For instance, a successful treatment of AML that develops from myelodysplastic syndrome (MDS) yields residual MDS clones that persist in the bone marrow (Masuya, Katayama et al. 2002, Batzios, Hayes et al. 2009).

Ultimately, functionalist oncology provides a framework for understanding context-dependent interactions. For example, it can explain how the addition of a SAMHD1 inhibitory drug interacts with ara-C treatment in AML, and how this interaction could influence the predicted anti-leukemic effect. Although substantial validation is required before any clinical implementation, in principle, this framework can be used to predict how the modeled outcomes will be affected by the frequency and sequence of a certain treatment. In this sense, this approach may help in taking a new step in the oncological decision-making process towards personalized treatment strategies.

## Code availability

The interactive simulation application AsymSAM is available as an R package at GitHub (private repository) and will be made publicly accessible upon publication: https://github.com/NaboulsiR/AsymSAM.

## Contributors

Conceptualization: AE, NH

Methodology: AE, ST

Investigation: ST, AE

Visualization: AE, ST, RN

Programming: RN, ST

Supervision: ST, MJ, NH

Writing—original draft: ST, AE, NH

Writing—review & editing: ST, AE, MJ, RN, NH

## Acknowledgments

Funding:

Swedish Children’s Cancer Foundation grants PR2020-0077 and TJ2019-0072 (NH)

Swedish Medical Association grant SLS-961737 (NH)

Swedish Research Council grant 2020-01184 (NH)

Radiumhemmet’s Research Foundations grant 191112 (NH)

Stockholm County Council grants K2892-2016 and 20200246 (NH)

Felix Mindus contribution to Leukemia Research grant 2019-01909

Karolinska Institutet Foundations grant 2-2109/2019-18

The authors thank Martin Sonntag and Vasilios Zachariadis for the scientific discussion.

## Supplementary Figures and Comments

To illustrate that the net benefit of pharmacologically targeting SAMHD1 depends on the parameters *x*, *y*, and *z*, we explicitly define several different AML clones with distinct *x*, *y*, and *z* and compare the total efficacy *e* ( *d* ) in the presence of a SAMHD1-inhibitory drug (Equation 3) with the efficacy of the ara-C treatment alone (Equation 2), as a function of *d* (Supplementary Figure S1).

**Supplementary Figure 1.**
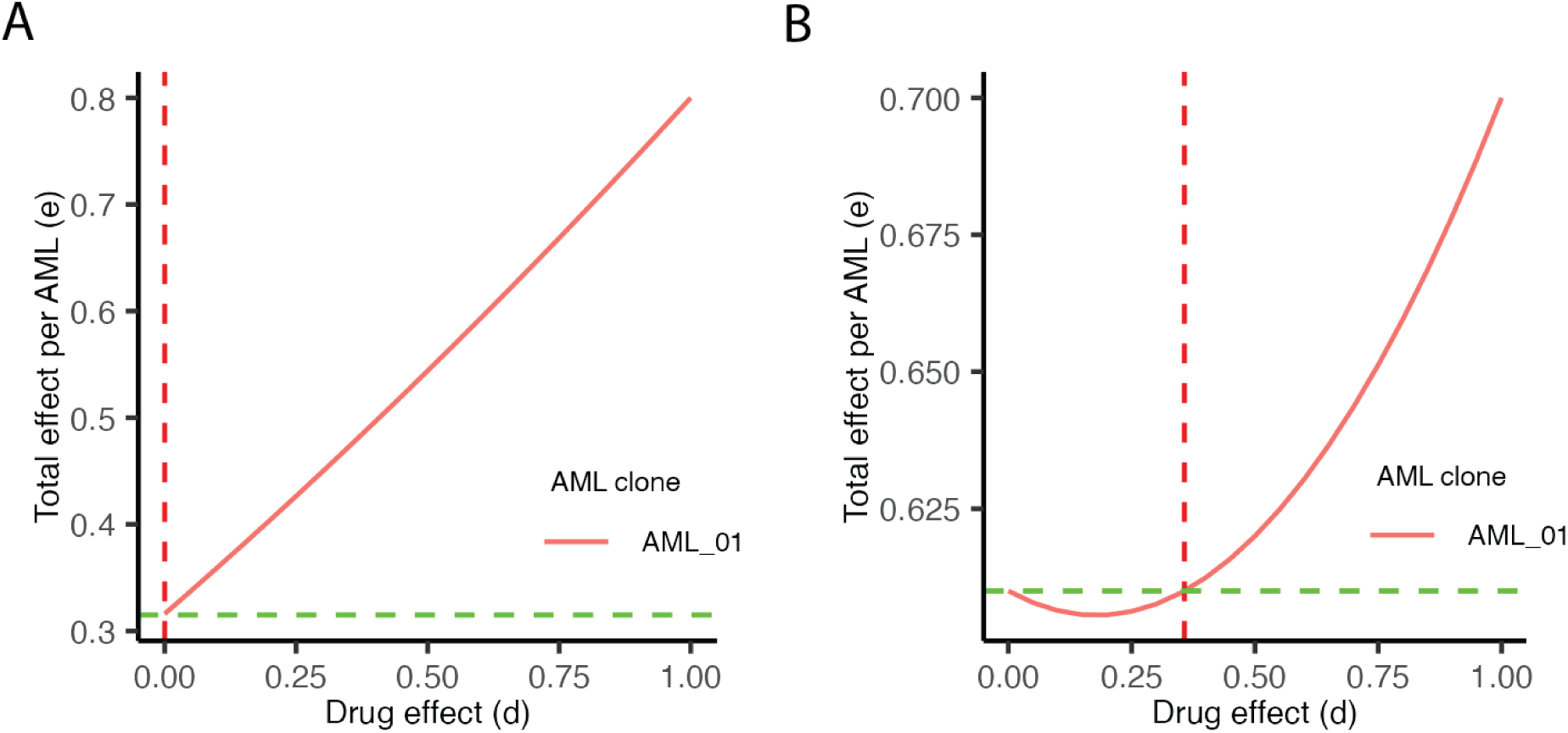
Total efficacy of the combined ara-C/anti-SAMHD1 treatment (y-axis) as a function of the efficacy d of the anti-SAMHD1 drug (x-axis). On all panels, the green dashed line (0.315 and 0.61, respectively) indicates the total efficacy without the anti-SAMHD1 drug, and the red dashed line (0 and 0.3571, respectively) indicates a threshold drug efficacy above which the anti-SAMHD1 drug is beneficial (whenever present). A) For AML1 defined with *x* = 0.8, *y* = 0.1, *z* = 0.7, administering the anti-SAMHD1 drug (orange curve) always has a net-positive effect on the chemotherapy. B) For AML1 defined with *x* = 0.7, *y* = 0.4, *z* = 0.5, inhibiting SAMHD1 has a net-positive effect on the chemotherapy (orange curve above green dashed line) only if *d* > 0.3571.

**Supplementary Table 1.**
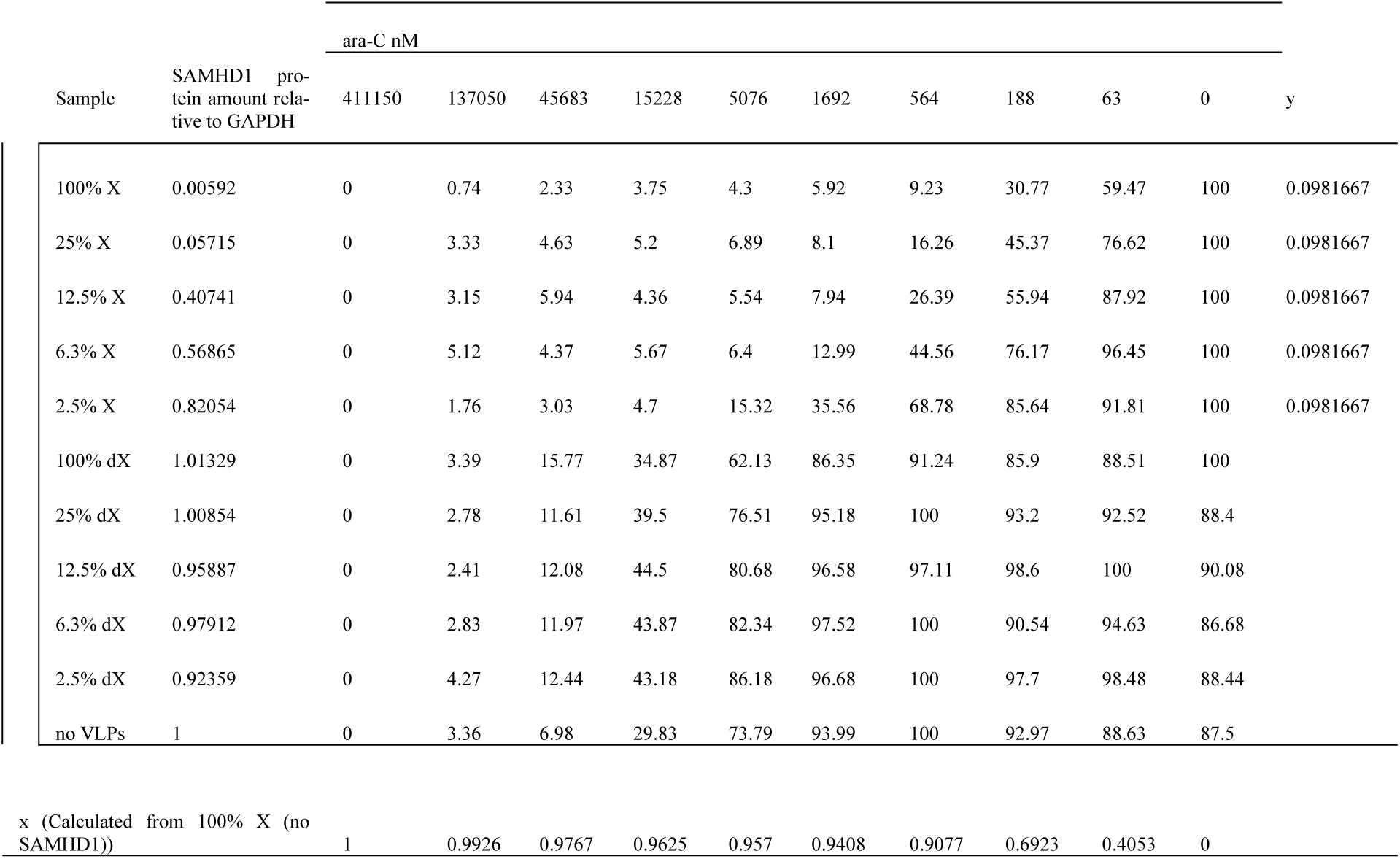
Experimentally derived parameters describing the Ara-C dose-response in THP-1 cells with variable SAMHD1 expression levels after treatment with Vpx-VLP (X), empty VLPs (dX), and no VLPs. The data present the various concentrations of Ara-C, their corresponding cell viabilities, the amount of SAMHD1 protein levels relative to GAPDH, and the derived model parameters: Ara-C efficacy (x), and SAMHD1 tumor-suppressive effect (y).

| Sample | SAMHD1 protein amount relative to GAPDH | ara-C nM |  |  |  |  |  |  |  |  |  |  |
| --- | --- | --- | --- | --- | --- | --- | --- | --- | --- | --- | --- | --- |
|  |  | 411150 | 137050 | 45683 | 15228 | 5076 | 1692 | 564 | 188 | 63 | 0 | y |
| 100% X | 0.00592 | 0 | 0.74 | 2.33 | 3.75 | 4.3 | 5.92 | 9.23 | 30.77 | 59.47 | 100 | 0.0981667 |
| 25% X | 0.05715 | 0 | 3.33 | 4.63 | 5.2 | 6.89 | 8.1 | 16.26 | 45.37 | 76.62 | 100 | 0.0981667 |
| 12.5% X | 0.40741 | 0 | 3.15 | 5.94 | 4.36 | 5.54 | 7.94 | 26.39 | 55.94 | 87.92 | 100 | 0.0981667 |
| 6.3% X | 0.56865 | 0 | 5.12 | 4.37 | 5.67 | 6.4 | 12.99 | 44.56 | 76.17 | 96.45 | 100 | 0.0981667 |
| 2.5% X | 0.82054 | 0 | 1.76 | 3.03 | 4.7 | 15.32 | 35.56 | 68.78 | 85.64 | 91.81 | 100 | 0.0981667 |
| 100% dX | 1.01329 | 0 | 3.39 | 15.77 | 34.87 | 62.13 | 86.35 | 91.24 | 85.9 | 88.51 | 100 |  |
| 25% dX | 1.00854 | 0 | 2.78 | 11.61 | 39.5 | 76.51 | 95.18 | 100 | 93.2 | 92.52 | 88.4 |  |
| 12.5% dX | 0.95887 | 0 | 2.41 | 12.08 | 44.5 | 80.68 | 96.58 | 97.11 | 98.6 | 100 | 90.08 |  |
| 6.3% dX | 0.97912 | 0 | 2.83 | 11.97 | 43.87 | 82.34 | 97.52 | 100 | 90.54 | 94.63 | 86.68 |  |
| 2.5% dX | 0.92359 | 0 | 4.27 | 12.44 | 43.18 | 86.18 | 96.68 | 100 | 97.7 | 98.48 | 88.44 |  |
| no VLPs | 1 | 0 | 3.36 | 6.98 | 29.83 | 73.79 | 93.99 | 100 | 92.97 | 88.63 | 87.5 |  |
| x (Calculated from 100% X (no SAMHD1)) |  | 1 | 0.9926 | 0.9767 | 0.9625 | 0.957 | 0.9408 | 0.9077 | 0.6923 | 0.4053 | 0 |  |

**Supplementary Table 2.** Experimental uncertainty calculations used for the estimation of the error propagation in the model.

| dX and no-VLP |  |  | de (uncertainty on e) | x | dx (uncertainty on x) | y | dy (uncertainty on y) | z | dz/de | dz/dx | dz/dy | dz (uncertainty on z) |
| --- | --- | --- | --- | --- | --- | --- | --- | --- | --- | --- | --- | --- |
| ara-C nM | Viability % | e total |  |  |  |  |  |  |  |  |  |  |
| 411150 | 0 | 1 | 0.094 | 1 | 0.094 | 0.09816667 | 0.049410674 | 0 | -1.10885234 | 1 | 0 | 0.140357881 |
| 137050 | 3.173 | 0.96826667 | 0.094 | 0.9926 | 0.094 | 0.09816667 | 0.049410674 | 0.02799474 | -1.11711902 | 0.97925172 | 0.03930872 | 0.139656157 |
| 45683 | 11.808 | 0.88191667 | 0.094 | 0.9767 | 0.094 | 0.09816667 | 0.049410674 | 0.11020475 | -1.13530494 | 0.91102206 | 0.1486534 | 0.137026861 |
| 15228 | 39.292 | 0.60708333 | 0.094 | 0.9625 | 0.094 | 0.09816667 | 0.049410674 | 0.41370033 | -1.15205438 | 0.60914252 | 0.50193461 | 0.124984489 |
| 5076 | 76.938 | 0.23061667 | 0.094 | 0.957 | 0.094 | 0.09816667 | 0.049410674 | 0.84653345 | -1.15867538 | 0.16036213 | 0.98850363 | 0.120313807 |
| 1692 | 94.383 | 0.05616667 | 0.094 | 0.9408 | 0.094 | 0.09816667 | 0.049410674 | 1.04950234 | -1.17862706 | -0.05261728 | 1.23351784 | 0.126545923 |
| 564 | 98.058 | 0.01941667 | 0.094 | 0.9077 | 0.094 | 0.09816667 | 0.049410674 | 1.09620152 | -1.22160663 | -0.10598383 | 1.32827991 | 0.132638117 |
| 188 | 93.152 | 0.06848333 | 0.094 | 0.6923 | 0.094 | 0.09816667 | 0.049410674 | 1.0475436 | -1.6016934 | -0.06867485 | 1.65441223 | 0.171441202 |
| 63 | 93.795 | 0.06205 | 0.094 | 0.4053 | 0.094 | 0.09816667 | 0.049410674 | 1.09881088 | -2.73588043 | -0.24379689 | 2.84544711 | 0.293989959 |
| 0 | 90.183 | 0.09816667 | 0.094 | 0 | 0.094 |  |  |  |  |  |  |  |

